# A Note on Computing Interval Overlap Statistics

**DOI:** 10.1101/517987

**Authors:** Shahab Sarmashghi, Vineet Bafna

## Abstract

We consider the following problem: Let *I* and *I*_*f*_ each describe a collection of *n* and *m* non-overlapping intervals on a line segment of finite length. Suppose that *k* of the *m* intervals of *I*_*f*_ are intersected by some interval(s) in *I*. Under the null hypothesis that intervals in *I* are randomly arranged w.r.t *I*_*f*_, what is the significance of this overlap? This is a natural abstraction of statistical questions that are ubiquitous in the post-genomic era. The interval collections represent annotations that reveal structural or functional regions of the genome, and overlap statistics can provide insight into the correlation between different structural and functional regions. However, the statistics of interval overlaps have not been systematically explored. In this manuscript, we formulate a statistical significance problem which considers the length and structure of intervals. We describe a combinatorial algorithm for a constrained interval overlap problem that can accurately compute very small *p*-values. We also propose a fast approximate method to facilitate problems consisted of very large number of intervals. These methods are all implemented in a tool, iStat. We applied iStat to simulated interval data to obtain precise estimates of low *p*-values, and characterize the performance of our methods. We also test iStat on real datasets from previous studies, and compare iStat results with the reported *p*-values using basic permutation or parametric tests. The iStat software is made publicly available on https://github.com/shahab-sarmashghi/ISTAT.git

## 1 Introduction

Annotating the genome is a central problem in biology. Subsequent to the sequencing and assembly of the human genome, and the development of deep sequencing technologies, researchers have employed creative ideas to develop better insight into the regions that support genome structure and function.

Examples of annotation include repeat elements [1], protein coding genes [2], non-coding RNA [3], regulatory regions [4], sites with specific epigenetic modifications [5], transcription start sites [4], ribosome initiation sites [6, 7]. Annotation may also involve structural features, such as the regions with a change in copy number and other structural variation [8, 9]. These regions can be dynamic and change depending upon tissue type and experimental conditions (e.g., histone methylation marks, regions with high gene expression, etc.), or relatively static (e.g., location of protein coding genes).

In all of these annotations, we can work with an abstract representation by considering the genome as a line segment, and any annotation as a collection of non-overlapping intervals on that line. Having a pair of annotations modeled by two sets of intervals, enables us to evaluate their overlap which is very useful in uncovering biological principles, and widely used by scientists.

Epigenetics is among the areas that extensively apply such models, in order to study the potential association between epigenetic modifications and functional elements in the genome. For instance, Guenther et al. 2007 [10] observed that about 3*/*4 of all known promoters were overlapped by the intervals highly enriched for the methylation of lysine 4 on histone H3, showing that a large fraction of genes are enriched for H3K4me3 modification, including genes without any detected transcript. Assuming that the presence of histone H3K4me3 is correlated with transcription initiation, they hypothesized that transcription initiation occurs in all genes, but only in active genes it is accompanied by transcriptional elongation.

This model can also be useful to study functional impact of segmental duplications and copy number variations (CNVs). Zarrei et al. 2015 [11] performed a meta-analysis and provided an updated map of CNVs in healthy individual. They have considered several sets of genes and genomic sequences such as protein-coding and non-coding genes, cancer genes, lincRNAs, Promoters, etc., and computed the enrichment of copy number variant regions in each of these annotations to assess the variability of different functional regions of the genome.

In experiments related to genome annotations, such questions are ubiquitous, and they all distill down to the underlying statistical question of significantly overlapping intervals. Hence, it has been a standard practice to formulate the problems as a hypothesis test and compute the *p*-value to substantiate the statistical significance of the observed overlap. Randomization tests are known to be exact significance tests when the space of all possible random samples can be enumerated. Random annotations can be generated by randomizing the position of intervals while preserving the coherence of each region. However, in many real-life examples including the above studies, the sample space is enormous, and naive sampling-based methods cannot achieve adequate resolution to distinguish between rare events in feasible running times. On the other hand, while parametric tests used in the literature are computationally efficient, they oversimplify the problem by casting intervals as points and ignoring the dimension of annotated regions on the genome, which often lead to more significant *p*-values.

In this paper, we propose an algorithm which efficiently enumerates over all possible randomized samples to find the exact null distribution, under the assumption that the order of intervals is preserved when randomizing their position. Using simulated data, we show that the impact of our assumption on *p*-value calculation is limited. We also provide a fast approximate solution based on Poisson binomial distribution, and using simulated data, we characterize its performance in approximating the generic null distribution. Moreover, we demonstrate the result of applying our methods to four examples of interval overlap problem from previously published studies, and compare our results with the *p*-values reported in these studies.

**Figure 1:**
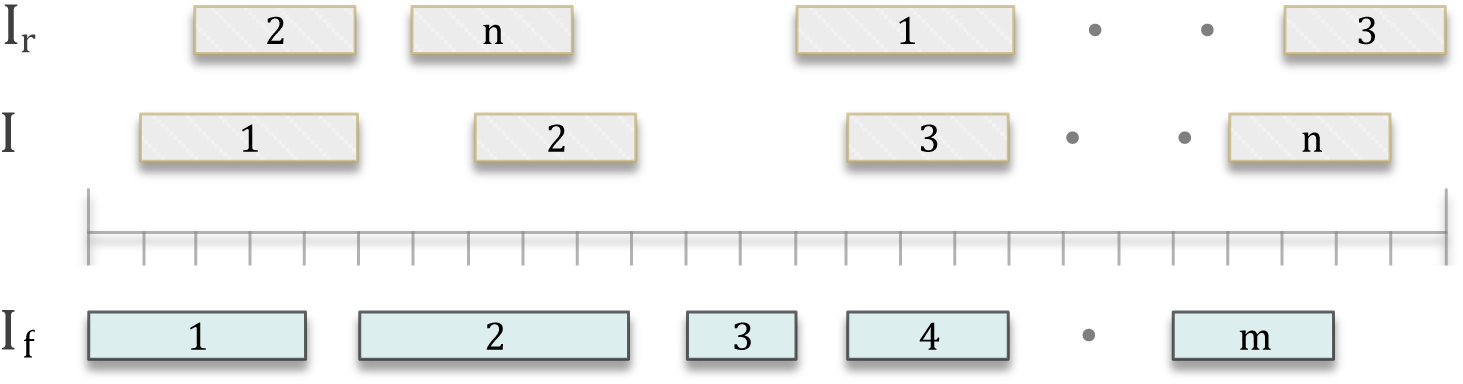
A Schematic of the interval overlap problem. *I*_*f*_ denotes the reference collection of intervals, and *I* represents the query collection. The randomized set *I*_*r*_ is generated by relocating the intervals in *I* such that all possible non-overlapping random sets are equiprobable.

## 2 Methods

Let us first introduce the notation frequently used throughout the paper. Let *I*_*f*_ denote a ‘reference’ collection of intervals, and *I* denote a ‘query’ collection of intervals (Figure 1). We use the space-counted, zero-start convention for the genomic coordinates. Namely, we count the space between bases starting from 0 (the one before the first base) up to *g* (the one after the last base), where *g* denotes the length of the genomic region of interest. Thus, each interval is denoted by a pair of indices (*u*_1_, *u*_2_) with 0 ≤ _1_ < *u*_2_ ≤ *g*, and is composed of the nucleotides between *u*_1_ and *u*_2_. We use ‘*i*’ to index the intervals in query set *I*, which has total number of *n* intervals, and designate ‘*j*’ to index the intervals in reference set *I*_*f*_, which consists of *m* intervals in total. The length of *i*-th query interval and *j*-th reference interval are represented by *l*_*i*_ and *x*_*j*_, respectively. Two intervals (*u*_1_, *u*_2_) and (*v*_1_, *v*_2_) overlap iff they share common nucleotide(s). A collection of intervals is *non-overlapping* if no pair of intervals in the collection overlap.

### Problem formulation

Let *I*_*f*_ ⊑*I* denote the subset of intervals in *I*_*f*_ that are *hit* (overlap with intervals in *I*). Suppose |*I*_*f*_ ⊑*I*| = *k*. We measure the significance (*p*-value) of this observation by sampling a random set of intervals *I*_*r*_ with the following properties (See Figure 1)

- |*I*_*r*_ = |*I*|. *I*_*r*_ has exactly *n* elements.
- Intervals in *I*_*r*_ have the same lengths as the intervals in *I*.
- The location of intervals in *I*_*r*_ are drawn from a distribution (implicitly) such that all possible random sets are equally likely.

Let *I*_*r*_ be drawn according to the process above, then *p*-value is defined as

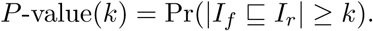

While the computational complexity of the problem is not known, we can argue that it is hard. Clearly, the number of possible random sets is very large; ranging from 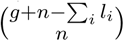 when all *l*_i_’s are identical, to 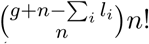 when all *l*_*i*_ are distinct. For typical values of *g* = 2 · 10^8^ (length of a chromosome), *n* = 100 (number of annotated regions), and Σ *l*_*i*_ = 10^6^ (total length of regions covered by an annotation), counting all possibilities naively to compute Pr(|*I*_*f*_ ⊑ *I* | ≥ *k*) is computationally intractable. Thus, we impose the restriction that the intervals in *I*_*r*_ must retain the same order as the intervals in *I*, and present a dynamic programming (DP) algorithm to compute the number of distinct random sets with |*I*_*f*_ ⊑*I*_*r*_| = *k*, for all *k*. In practice, to apply the algorithm to large genomes with abundant annotation we use a practical interval ‘scaling’ scheme by considering the natural partitioning of the genome into intervals and the gaps amidst them, and scale each interval and gap in *I* and *I*_*f*_ by a fraction *ν*. Ideally, we want to have *ν* = 1, but large problems require smaller fractions to make the computation feasible from both running time and memory usage aspects. Nevertheless, we show that the algorithm still yields a close approximation of *p*-value.

### 2.1 Dynamic programming algorithm

For interval *i* in *I*_*r*_, genomic location *h*, (1 ≤ *h* ≤ *g*), 0 ≤ k ≤ *m*, *a* ∈ 0, 1, let *N* (*i, h, k, a*) denote the number of arrangements of the first *i* intervals in *I*_*r*_ such that (see Figure 2):

- The *i*-th interval ends exactly at location *h*.
- *k* intervals in *I*_*f*_ are hit by the first *i* intervals in *I*_*r*_.
- *a* = 0 if the interval from *I*_*f*_ that spans *h* (if any) has not been counted earlier; *a* = 1 otherwise.

**Figure 2:**
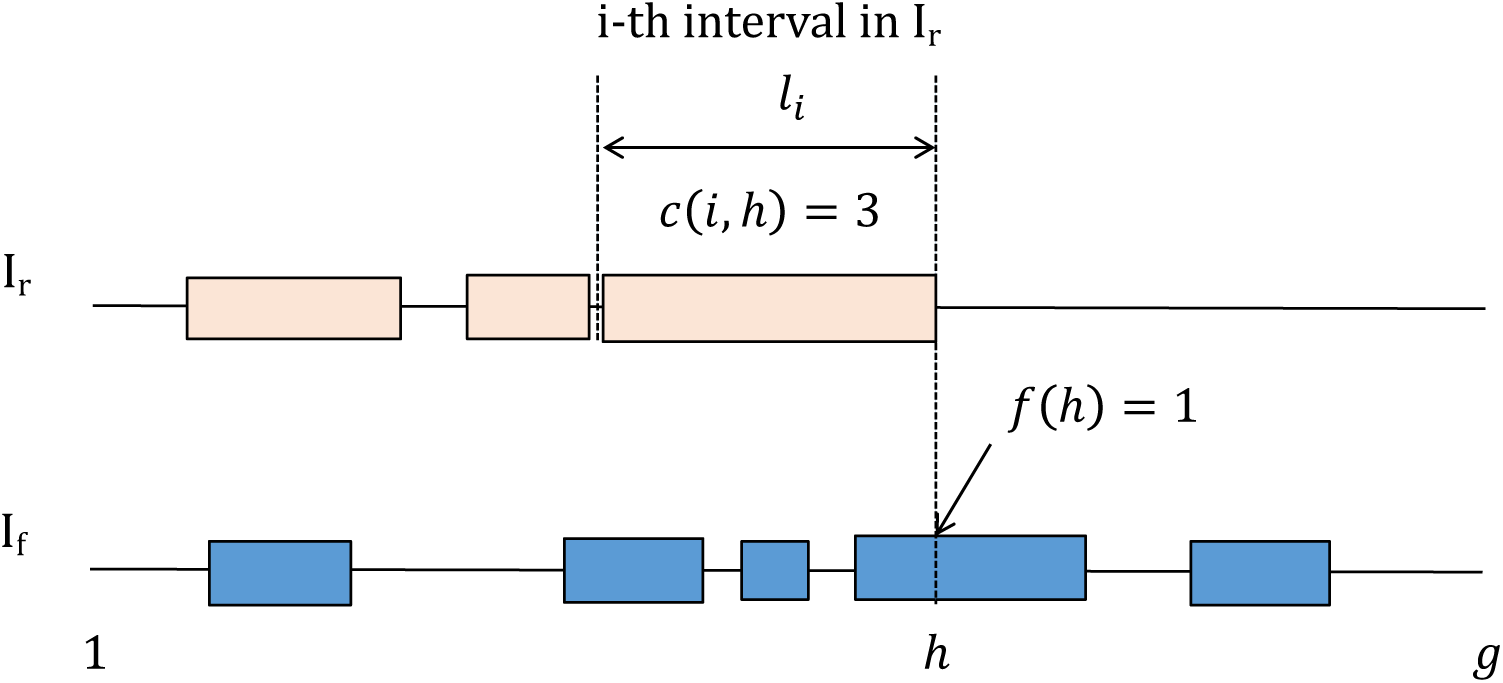
Cartoon for Dynamic Programming

We also define *N*_1_(*i, h, k, a*) identically to *N* (*i, h, k, a*) with the exception that the *i*-th interval ends at or before location *h*. Note that if the *j*-th interval in *I*_*f*_ spans *h*, it is counted as a hit, but may have already been counted by some other interval in *I*_*r*_. Although a separate function can be defined to store that information, we use *a* as an indicator in dynamic programming for the sake of brevity. In order to compute *N*_1_(*i, h, k, a*), we must define some auxiliary functions. Let *c*(*i, h*) denote the number of intervals in *I*_*f*_ which intersect with (*h* – *l*_*i*_, *h*) in *I*_*r*_. While evaluating *c*(*i, h*), (*j*_1_, *j*_2_) in *I*_*f*_ is counted as an intersecting interval with (*h − l_i_, h*) if *j*_1_ *< h* and *j*_2_ *> h −l_i_*. We also define binary function *f*: (0*, g*] → {0,1}, where *f* (*h*) = 1 if some interval in *I*_*f*_ spans *h*, and *f* (*h*) = 0 otherwise. See Figure 2. For the simplicity of exposition, it is assumed that a single nucleotide overlap between two intervals from *I_r_* and *I*_*f*_ is sufficient to count the reference interval as intersected. The generalization of algorithm to accommodate stricter conditions is straightforward and can be done by modifying the definition of *c*(*i, h*) and *f* (*h*) (Appendix 1.1).

To explain the recurrences, note that *N*_1_(*i, h, k, a*) can be computed by adding cases where the *i*-th interval ends exactly at *h*, and cases where the *i*-th interval ends strictly before *h*. To compute *N* (*i, h, k, a*) we need to consider all arrangements where the first *i −*1 intervals in *I_r_* ends before the start of the *i*-th interval at *h −l_i_*.

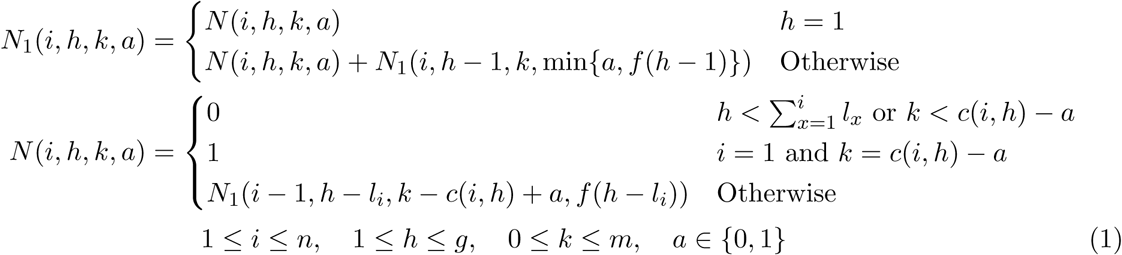

We note a technical difference between non-overlapping and ‘disjoint’. Intervals (*i*_1_, *i*_2_) and (*i*_2_, *i*_3_) are non-overlapping as they do not share any nucleotide, but are not disjoint because we cannot distinguish them from interval (*i*_1_, *i*_3_). The case where *I*_*r*_ is restricted to be disjoint is described in Appendix 1.2. The *DP p*-value (Pr(|*I*_*f*_ ⊑*I*_r_| ≥ *k*)) can be computed using the ratio

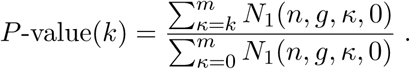

Recall that the total number of configurations is

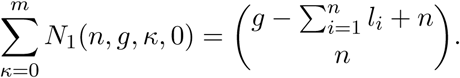

which can be very large and surpass the upper limit of ordinary data types. Therefore, we perform all calculations using a logarithmic scale (Appendix 1.3).

#### Time complexity

The number of iterations to complete the table of values for *N*_1_(*i, h, k, a*) is 𝒪(*ngm*). The functions *c*(*i, h*) and *f* (*h*) can be pre-computed (using a modified version of binary search algorithm), so each iteration is computed in a constant time. Therefore, the total time complexity is 𝒪(*ngm*) which is pseudo-polynomial because the input size is 𝒪((*n* + *m*) log *g*). The running time can be reduced to 𝒪(*ngνm*) by scaling the genome using scaling factor *ν <* 1. We also use a number of tricks to improve the speed of computations, including lowering memory usage from 𝒪(*ngm*) to 𝒪(*gm*). We should note that this time complexity is achieved under the assumption that the order of intervals in *I*_*r*_ is same as *I*. In Results, we show that choosing different orders does not significantly change the *p*-value.

#### Multiple chromosomes

In many cases of interest, the intervals reported are on multiple chromosomes, with a non-uniform distribution across chromosomes. Therefore, the appropriate random interval set 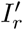 may only allow permutation of interval positions within the chromosome it is originally assigned to. For this alternative null model, the DP algorithm is applied to each chromosome to enumerate rearrangements of intervals within each chromosome, and then the results are combined to compute the overall *p*-value. See Appendix 1.4.

### 2.2 Poisson binomial approximation

For the case that annotations contain too many intervals such that the processing resources to run DP algorithm cannot be afforded, we provide an approximation which is reasonably close under certain condition. For simplicity, we remove the non-overlapping assumption on *I*_*r*_. Thus, *I*_*r*_ is a randomly located collection of *n* intervals of lengths *l*_1_, *l*_2_, *l*_3_, …, *l*_n_ with arbitrary order. Let *E*_*ij*_ denote the event that the *j*-th interval in *I*_*f*_ is intersected by the *i*-th interval in *I*_*r*_. Then,

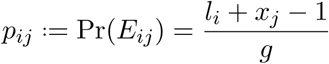

As before, we assumed that a single nucleotide overlap is sufficient, but it can be easily generalized to a more strict overlap condition (Appendix 1.1). Let 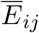 be the event that the *i*-th interval in *I_r_* does not intersect the *j*-th interval in *I*_*f*_. In the absence of the non-overlapping assumption on *I*_*r*_, the events 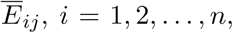 are independent, and the probability of their intersection is given by the product of individual probabilities. Therefore, the probability of 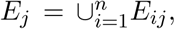, which is the event where interval *j ∈ I_f_* is hit by *I_r_*, can be calculated as

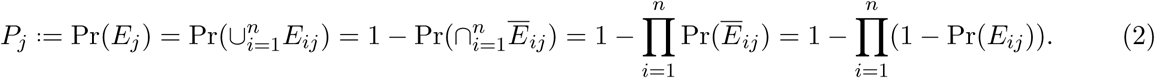

**Figure 3:**
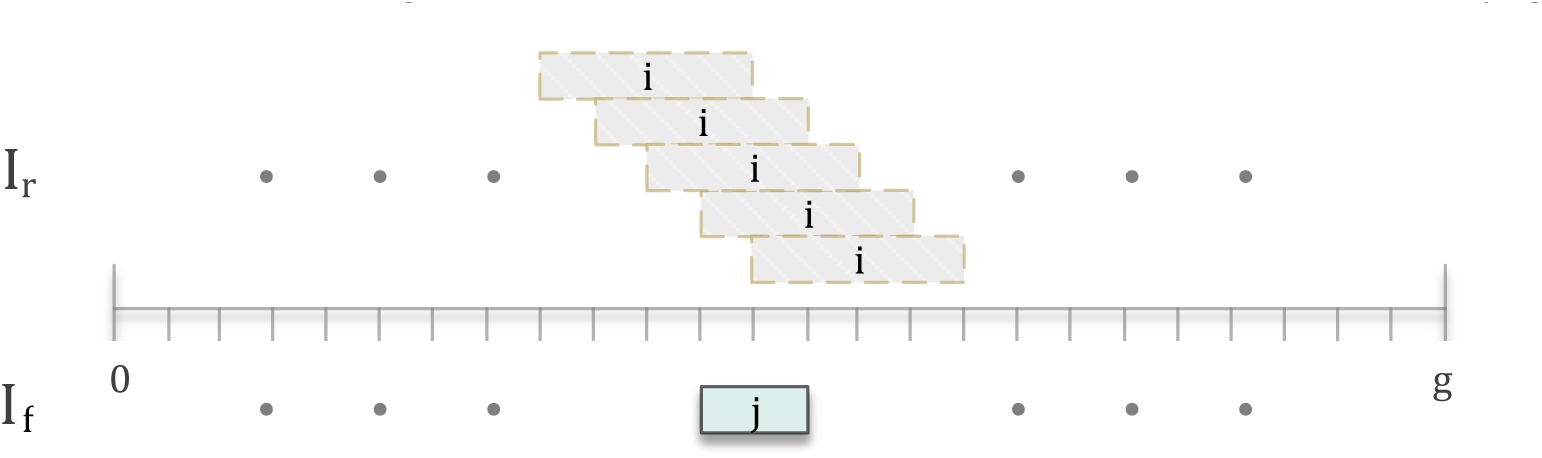
Illustration of the cases that *i*-th interval from *I_r_* intersects *j*-th interval in *I*_f_

Now consider the binary indicator variable *X_j_*, where *X_j_* = 1 iff event *E_j_* occurs. We have *m* Bernoulli experiments with success probabilities *P*_1_*, P*_2_*, …, P_m_*, and we are interested in computing Pr(Σ_*j*_*X*_*j*_ = *k*). In general, there are dependencies between *E*_*j*_’s for different values of *j*. However, under certain condition where intervals are not too close or far away, we can approximately assume independence between different intervals. The sum of *m* independent Bernoulli trials with different success probabilities is a Poisson binomial (PB) distribution [12].

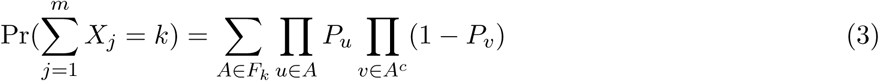

where *F*_*k*_ is the set of all subsets of {1, 2, …, *m*} with *k* elements. Eqn. 3) allows us to compute the *p*-value as

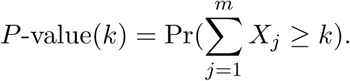

We cannot directly use Eqn. 3 by enumerating over all elements in *F_k_*, but use a recursive approach to compute it, following Hong [13]. It is reproduced here for completeness. Let 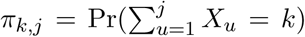 denote the probability of getting *k* hits in the first *j* intervals in *I*_*f*_. Our goal is to compute 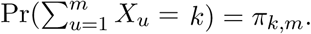 All values π_*k;j*_ can be computed in 𝒪(*m*^2^) time using

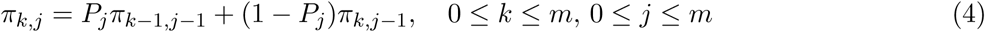

with the boundary conditions *π*_–1, *j*_ = *π*_*j*+1*, j*_ = 0, *j* = 0, 1, …, *m* and *π*_0,0_ = 1. Other FFT based methods are also applicable [13].

With the above PB approximation, we assume that the event of an interval in *I*_*f*_ being hit is independent of other intervals being hit, greatly reducing the computational complexity of the problem. To understand the impact of this assumption, we introduce a new parameter. Recall that *P*_*j*_ = Pr(*E*_*j*_) = Pr(*X*_*j*_ = 1) is the probability that interval *j* (length *x*_*j*_) in *I*_*f*_ is hit by some interval in *I*_*r*_. Let *d*_*j*_ denote the distance of interval *j* from interval *j* –1. Define ∆≔ (*m* –1). **median**. {*d*_*j*_|*j* = 2,3, …, *m*}, and 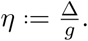. Parameter *η* is a measure of the ‘spread’ of intervals in *I*_*f*_. For *η ≪*1, and *j’* sufficiently close to *j*, we expect to have

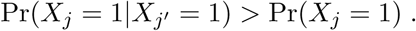

In other words, if intervals in *I*_*f*_ are clumped, then 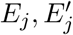 are not statistically independent but positively correlated, and we will underestimate the true *p*-value. For larger values of *η*, and *j, j’* sufficiently distant,

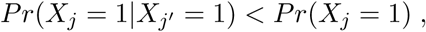

The negative correlation leads to an over-estimation of the *p*-value. To better recognize this effect, imagine an extreme case where *n < m* and due to the size and spread of intervals in *I*_*f*_, at most *n* intervals in *I*_*f*_ can be hit. Therefore, *p*-value(*n* + 1) = Pr(Σ_*j*_*X*_*j*_ > *n*) = 0. The independence assumption in PB computation, though, will lead to a non-zero value (over-estimate) for *p*-value(*n* + 1).

## 3 Results

### 3.1 Performance on simulated data

We simulated intervals in a randomly generated chromosome to test the performance of iStat. To study the impact of scaling and fixed-order assumption on the DP algorithm, we chose *g* = 200Mbp, and the two sets of intervals *I* and *I*_*f*_ with *n* = *m* = 100 intervals. The intervals in *I* and *I*_*f*_ are generated with random lengths *l_i_* and *x*_*j*_ distributed uniformly over [1Kbp, 10Kbp]. The intervals in *I*_*f*_ were placed uniformly at random along the chromosome, while ensuring no overlap between them. To benchmark the speed of DP algorithm, we changed *n*, *m*, and *g* over a range of values and measured the running time of iStat. We also simulated intervals in *I*_*f*_ distibuted non-uniformly over the chromosome to study how their positional distribution impacts the quality of PB approximation.

#### The impact of scaling on DP *p*-value

The algorithm has substantial demands on memory and time. To allow it to work on the human genome, we scale down the intervals and the gaps between them by a fraction *ν*. To test the impact of scaling, we considered the example of a chromosome described above, with *g* = 200Mbp, and *n* = *m* = 100. The impact on DP *p*-values due to scaling with *ν ∈* {1,10^−1^, 10^−2^, 10^−3^} is shown in Figure 4a. As can be observed, scaling preserves the *p*-values tightly. To further investigate robustness of DP *p*-value computation to the scaling, we also considered an adversarial example where *I* and *I*_*f*_ contain intervals smaller than *ν^−1^*. For that purpose, the length of intervals generated from a uniform distribution over [100bp, 4Kbp], thus when we apply scaling factor *ν* = 10^−1^ about one fourth of intervals are smaller than the resolution *ν*^−1^ and become unit intervals. Nevertheless, *p*-values obtained with *ν* = 10^−1^ tightly followed finer-scale *p*-values (Figure S1a), verifying that we can apply reasonable scaling factor (with respect to the distribution of the length of intervals and gaps) to facilitate *p*-value computation using DP algorithm.

#### Effect of order on *p*-value

To test the effect of fixed order on *p*-value, we used a scaling factor *ν* = 10^−1^, and applied the DP method to 100 random instances of simulated intervals described before, each with a random permutation of *I*. In Figure 4b, we plotted the mean *p*-value for all *k*, as well as the standard error of the mean. We observe that the standard error is distributed tightly around the mean, while its ratio to the mean increases slightly for smaller *p*-values. The mean *p*-value range from 0.4320±2.279·10^−3^ for *k* = 1 to 1.017 · 10^−269^ ±6.246 ·10^−7^ for *k* = 100. The results suggest that fixing the order in DP algorithm to compute the *p*-value is an acceptable compromise for many real data-sets.

#### Running time

Using a desktop PC with Intel Core i7-6700K CPU and 32GB DDR4 RAM, the running time of our DP algorithm (in a logarithmic scale) versus the number of query intervals is plotted in Figure 4c for a number of scaling factors. The running time scales almost linearly with the number of query intervals *n*. It also scales linearly with the number of reference intervals *m* (Figure S1b) and the size of chromosome *g* (Figure S1c), and when larger scaling factors is used.

#### PB versus DP

To test the role of *η* in *p*-value estimation, we compared the *p*-values of the Poisson binomial method against the DP method for different values of *η*. See Figure 4d–f. Relative to the DP, the PB approximation underestimates *p*-values when *η* = 0.005445 (Figure 4d), and over-estimates for *η* = 0.6197 (Figure 4f). However, this over-estimation is not as pronounced as the under-estimation in the case of clumping, and reduces with large *n* (Figure S1ef). In our simulations, we changed the number of query and reference intervals as well as the spread of reference intervals over the genome, and compared the p-values computed using each method. Although the closeness of PB approximation is a complicated function of the distribution of intervals and its exact characterization is hard, as a rule of thumb we suggest to consider using DP (with the largest computationally-feasible scaling *ν*) when *η <* 0.06, to avoid liberal *p*-values (more significant). In the case that we have multiple chromosomes, the minimum *η* among all chromosomes can be considered to be as conservative as possible.

### 3.2 Enrichment analysis on real data

We also took four examples from the literature and applied iStat to to test its performance on interval data from previously published studies and compare the p-values estimated by iStat with the reported p-values. The first example comes from [14], relating to matching of focal copy number changes in tumor genomes. The second dataset is from [11] where a map of copy number variation (CNV) in the human genome is provided, and different genomic elements are investigated for the presence/absence of CNVs. We also ran iStat on an example from epigenetics context [10], where the promoters are found to be enriched for H3K4 methylation. The last example was extracted from an effort to systematically annotate genome by the means of characterizing chromatin states [15].

#### TCGA-CNV enrichment in HIRT (extra-chromosomal data)

For *I*_*f*_, we chose a collection of intervals with recurrent copy number amplifications in the TCGA array CGH data-set (named TCGA-CNV) [16, 17]. For *I*, we used amplified genomic regions from a whole genome sequencing experiment with an experimental protocol, HIRT, that preferentially selected extra-chromosomal fragments. A strong enrichment of TCGA-CNV intervals in the HIRT intervals would suggest that *many copy number amplifications can be attributed to the formation and independent replication of episomes (extra-chromosomal elements)*. The number of intervals in query and reference sets were not large, *n* = 116 and *m* = 101, so we did not scale the intervals and the resulting *p*-value is 8.679 10–6. For comparison, we applied scaling factor *ν* = 10^−1^ and the change in *p*-value was very small. As expected from *η* = 0.001, PB approximation gives more significant *p*-value = 2.642 ·10^−10^ (Figure 5a).

#### Non-coding genes enrichment in CNVs

We chose the set of all CNV gains from the inclusive map as *I*, and the set of all non-coding genes as *I*_*f*_, containing *n* = 3132 and *m* = 9058 intervals, respectively. Using the scaling factor *ν* = 10^−2^, we obtained *p*-value = 5.216 · 10^−18^, confirming high enrichment of non-coding genes in CNV gains. After applying an order of magnitude smaller scaling factor *ν* = 10^−3^, we get very close *p*-value = 2.532 ·10^−18^ which shows that scaling with *ν* = 10^−2^ is fine (Figure 5b). PB approximation *p*-value is 1.370 ·10^−52^, much smaller *p*-value that is consistent with *η* = 0.024. In the paper, they consider the exons of non-coding genes as *I*_*f*_, and report *p*-value = 0.0001 from a 10000 randomized dataset, which shows the limited resolution of basic permutation tests. The result of our algorithm indicates that computing the exact *p*-value in this case requires at least about 10^18^ randomized samples, which is impossible. In the supplementary they have also reported a binomial *p*-value = 2.32 ·10^−54^.

#### Enrichment of H3K4me3 in promoters

In [10] authors found that 74% of all annotated promoters were enriched for H3K4 methylation, concluding that a large fraction of genes with no detected transcript have promoter-proximal nucleosomes enriched for H3K4me3 modification. To evaluate the statistical significance of this observation, we took the set of regions highly enriched for H3K4me3 in ES cells (provided as supplementary information in [10]) as the query set. However, they have not provided the coordinates of promotors, and so for the reference intervals, we used the collection of all promoters (–5.5Kbp to 2.5Kbp relative to TSS of all RefSeq genes) as the reference set of intervals. Although with *I*_*f*_ that we used we did not get the same ratio of overlap as reported in the paper, but still the p-value is quite significant. At the observed overlap, PB *p*-value is 1.775 ·10^−76^, while DP *p*-value with *ν* = 10^−2^ is 2.734 ·10^−82^. For this example, *η* = 0.1 so PB approximation gives conservative *p*-values as expected (Figure 5c).

#### Enrichment of promoters in promoter-associated chromatin states

Among 51 identified chromatin states, states 1 to 11 were referred to as promoter-associated states because of high enrichment for promoter regions. We tried to compute the *p*-value of enrichment by considering the set of all promoter regions (within 2Kbp of RefSeq TSS) as the query set *I*, and 200-bp intervals identified with state 9 as the reference set *I*_*f*_. The *p*-value = 1.588 ·10^−8^ (under the scaling factor *ν* = 10^−2^) shows that it would be very unlikely to observe such overlap only by chance, yet it is much less significant than the *p*-value reported by the authors (≤10^−200^), computed using hypergeometric distribution. As *η* = 0.01, PB approximation expectedly gives liberal *p*-value (Figure 5d).

## 4 Discussion

Our results explore the statistics of interval overlaps. The question is quite natural in the post genomic era where annotating the genome for function, structure, and variation and identifying correlated annotations is a key problem. While scientists have used many different ways to compute the significance of overlap between two sets of intervals, their computations often do not explicitly state the assumptions on the null model, or accurately compute the *p*-values given specific assumptions.

To the best of our knowledge, the *p*-value computation for sets of overlapping intervals has been limited to either permutation tests which do not scale to computation of small *p*-values, or simple parametric tests such as hypergeometric or binomial tests which are based on simplifying assumptions about the length and structure of intervals. Our paper, however, formulates a null model where the size of intervals and their relative arrangement are considered when the significance of overlap is evaluated. We explicitly state the assumptions that we have made in our proposed model, and assess the impact of our assumptions thorough the experiments on simulated and real datasets. Computation of exact *p*-values may be necessary in some cases. For example, *p*-values can be used to compare the significance of two ‘competing’ annotations with different numbers of intervals (*n*) and intersections (*k*). We develop a novel frame-work that makes exact computation of *p*-value possible, even for very small *p*-values.

The proposed DP method is able to compute very small *p*-values by efficiently counting the number of possible random rearrangements of intervals resulting in specific amount of overlap. Although we assume that the order of intervals is not changed, and it may be possible to construct adversarial examples where changing the order has a material impact on *p*-values, but our simulation of typical examples of interval data show that the resulting change in *p*-values is not significant. Our experiments on simulated and real datasets also suggest that to improve the speed and memory usage, we can employ reasonable scaling factors and sill obtain accurate *p*-values.

The Poisson binomial approximation is very efficient to compute. However, our results suggest that for typical values found in real-life examples, the independence assumption is too strong, and might result in under-estimated *p*-values, or the false reporting of some overlap as being significant. Nevertheless, we have introduced parameter *η* which can be readily computed from the data before running the DP method, to estimate the accuracy of PB method compared to DP algorithm results. Future work should look into more systematic characterization of PB approximation.

Throughout our experiments, we let the intervals to be uniformly distributed over the whole extent of chromosomes. However, one might be interested in a non-uniform distribution of intervals under the null model, to account for confounding variables such G/C content, sequence context, or intergenic/genic region. Our methods can be used in such cases by confining the problem to the specific regions of interest. Hence, only intervals falling into such regions are considered, and *g* would be the total length of the segments that intervals are allowed to be distributed there. Moreover, we considered the overlap of two intervals as a binary event, and defined the statistic based on the number of overlapping intervals. However, DP method can be modified to compute the *p*-value when the overlap statistic is defined based on the total amount of shared base pairs instead. Thus, we provide this as an option in iStat software and give the user the flexibility of choosing the appropriate measure of overlap for their specific application.

**Figure 4:**
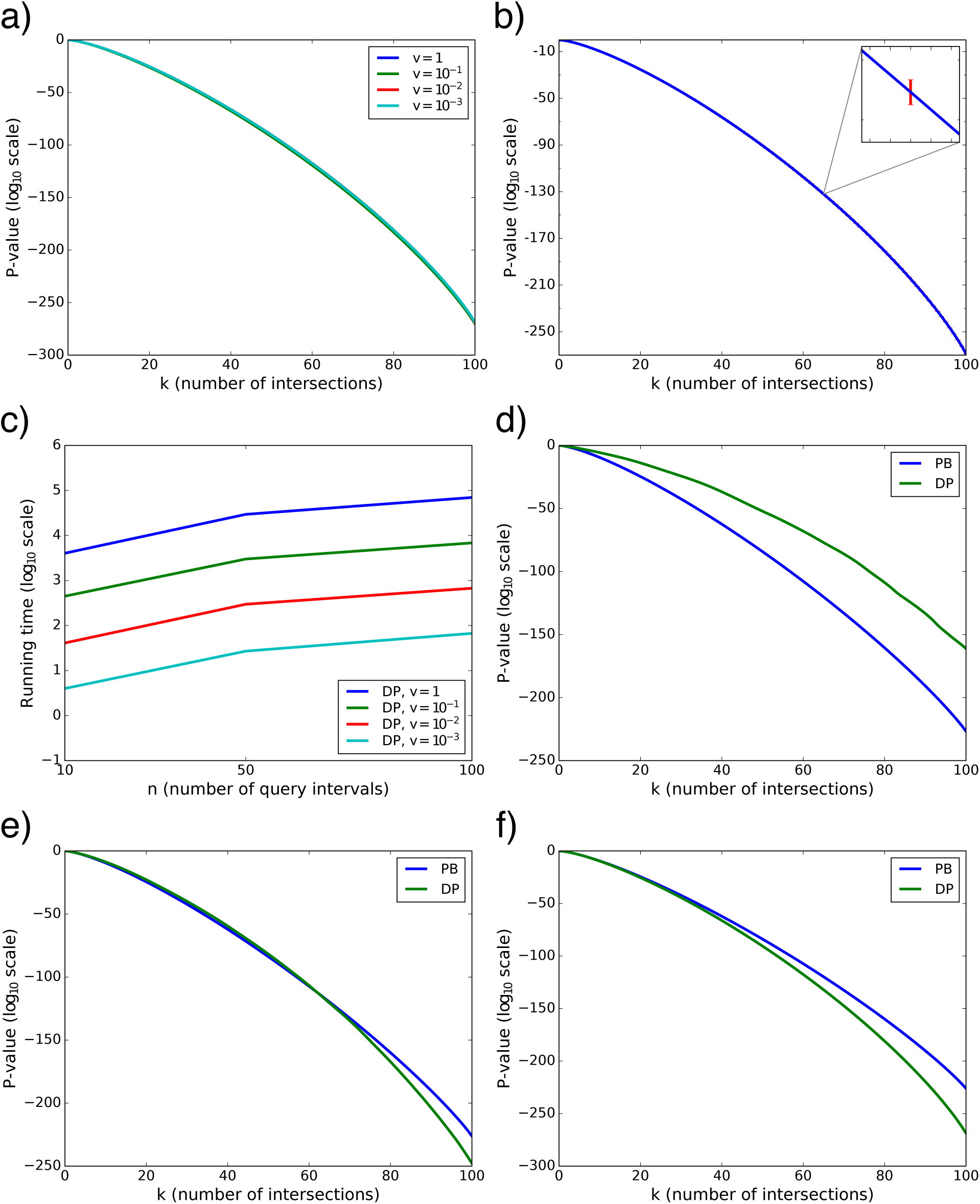
Testing methods on simulated data. (a) Impact of scaling parameter *ν* on DP *p*-value when *l_i_, x_j_*𝒖[1Kbp, 10Kbp]. (b) Impact of ordering on DP *p*-value, with *ν* = 10^*−*3^. The mean of 100 *p*-value computations for random orderings is plotted, and the error bars represent the standard error of the mean. (c) Running time (in secs.) of DP algorithm as a function of *n*, with *m* = 100 and *g* = 200Mbp. (d–f) Impact of approximation on *p*-value computation. Simulations are run with *g* = 200Mbp, *m* = 100, *n* = 100, *l*_*i*_, *x*_*j*_ [1Kbp, 10Kbp]; (d) *η* = 0.0054, (e) *η* = 0.053, (f) *η* = 0.62.

**Figure 5:**
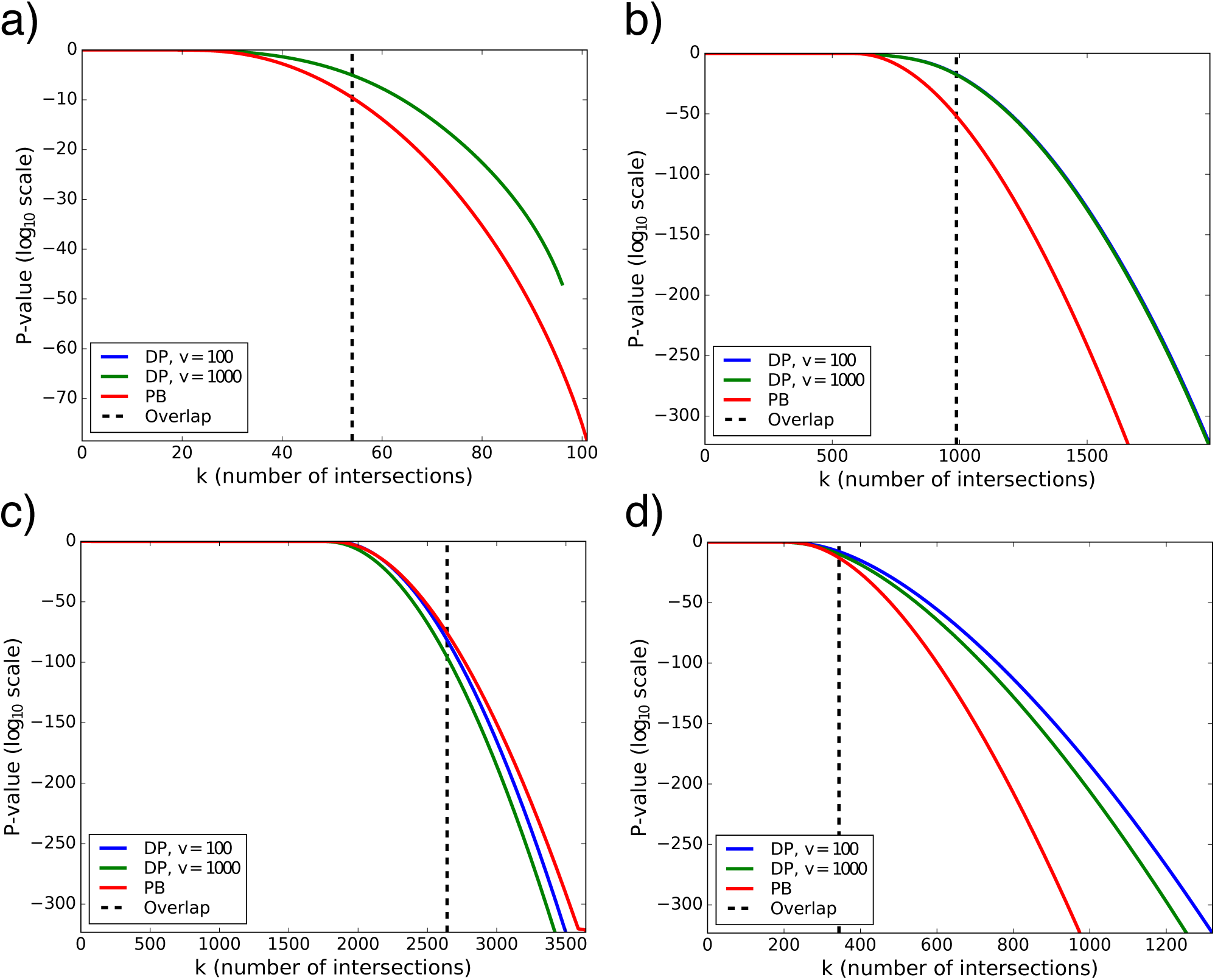
Enrichment analysis on real datasets. (a) TCGA-CNV enrichment in HIRT (extra-chromosomalcdata). (b) Non-coding genes enrichment in CNVs. (c) Enrichment of H3K4me3 in promoters. (d) Enrichment of promoters in promoter-associated chromatin states.

## 1 Appendix

### 1.1 Generalized overlap

We can be more strict about declaring an intersection by accepting only those overlaps which include *z* or more base pairs (units). The dynammic programming algorithm and Poisson binomial approximation can both be easily generalized for that:

#### 1.1.1 DP algorithm

For *c*(*i, h*), the intersection conditions should change to *j*_1_ ≤h –z and *j*_2_ ≥ *h* –*l*_*i*_ + *z*, which can be compressed into the single condition min {*j*_2_, *h*}–max {*j*_1_, *h* − *l*_*i*_} ≥ *z*. For *f* (*h*) we need to modify the defintion of “span”, so interval (*j*_1_, *j*_2_) spans *h* if *j*_1_ ≤ h –*z* and *j*_2_ ≥*h* + *z*, which allows it to have the opportunity of overlap (under this new criteria) with both intervals starting and ending at *h*.

#### 1.1.2 Poisson binomial approximation

In this case, *p*_*ij*_ is given by

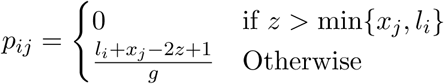

### 1.2 Dynamic programing with disjointed Intervals

As described earlier, for *i*-th interval in *I*_*r*_, genomic location *h*, (1 ≤*h* ≤g), 0 ≤*k* ≤*m*, *a* ∈0, 1, let *N* (*i, h, k, a*) denote the number of arrangements of the first *i* intervals in *I*_*r*_ such that (See Figure 2):

- The *i*-th interval ends exactly at location *h*.
- *k* intervals in *I*_*f*_ are hit by the first *i* intervals in *I*_*r*_.
- *a* = 0 if the interval from *I*_*f*_ that spans *j* (if any) has not been counted earlier; *a* = 1 otherwise.

We also define *N*_1_(*i, h, k, a*) identically to *N* (*i, h, k, a*) with the exception that the *i*-th interval ends at or before location *h*. If we consider the restriction that the intervals in *I*_*r*_ must be disjoint, which means that for any ordered pair of intervals (*i*_1_, *i*_2_) and (*i*_3_, *i*_4_), *i*_2_ has to be strictly less than *i*_3_, then the recurrence relation for *N* (*i, h, k, a*) has to be modified as:

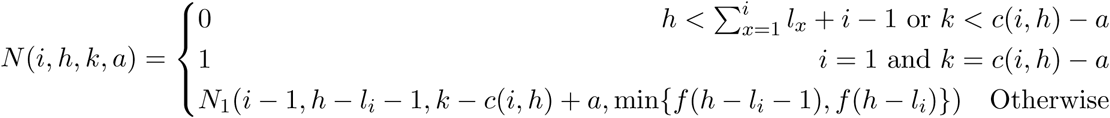

### 1.3 Log-scale computations

Let *a* = log *A* and *b* = log *B*, then the following simple math trick enables us to calculate *c* = log(*A ± B*) without explicitly converting *a* and *b* to their intractably large counterparts *A* = exp(*a*) and *B* = exp(*b*)

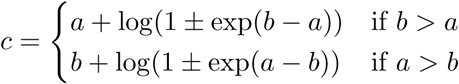

As a matter of fact, this trick is useful when *A* and *B* are both large, but the ratio 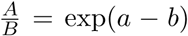 is computable, which is the case in the recurrence relation given by Eqn. 2.1. In fact, as we proceed along the four dimensions of *N*_1_(*.,.,.,.*), configurations accumulate and their count increases gradually. Therefore, whenever two numbers are added, their ratio is within the admissible range, even if their absolute values are not. The multiplication and division can be also done trivially.

### 1.4 The null model with multiple chromosomes

Consider *Q* chromosomes. For arbitrary chromosome *q*, let *I*_*q*_ ⊆*I* and *I*_*f, q*_ ⊆*I*_*f*_ denote the subsets of intervals paced on *q*, containing *n*_*q*_ and *m*_*q*_ intervals, respectively. Similarly, we can define *I*_*r, q*_ to be a random reordering of *I*_*q*_ on chromosome *q*. Let *N*_*q*_(*k*_*q*_) denote the number of configurations of intervals in *I*_*r, q*_ s.t. |*I*_*f, q*_ ⊑*I*_*r, q*_| = *k*_*q*_. Using dynamic programming on each of *Q* chromosomes, we can obtain *N*_*q*_(*k*_*q*_) 1 ≤ *q* ≤ *Q*, 0 ≤*k*_*q*_ ≤*m*_*q*_. For *k* ∈[0, *m*] we define the *p*-value to be

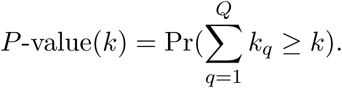

With the equiprobability assumption and using simple arguments based on multiplication principle to count the number of desired configurations, we can compute the *p*-value as

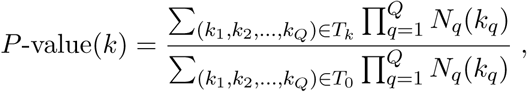

where *T*_*k*_ is the set of all *Q*-tuples (*k*_1_, *k*_2_, …, *k*_*Q*_) such that 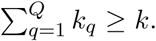 While the denominator can be easily computed via the following identity

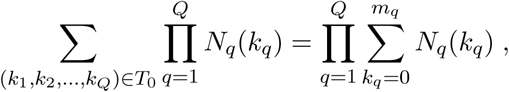

it is not efficient to iterate over *T*_*k*_ to compute the numerator for each *k*. Instead, we use a simple recursive procedure to compute it. Let *M* (*q, k*) be the number of configurations that the first *q* chromosomes have *k* intersections. The *p*-value can be expressed in terms of *M* (*q, k*) as

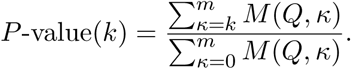

The following recurrence relation lets us to efficiently compute the *p*-value for all *k ∈*[0, *m*]

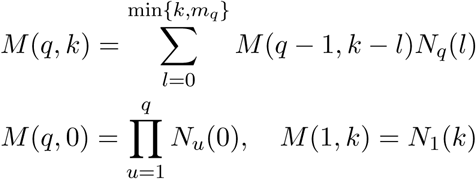

where the time complexity is 𝒪(*Qm*^2^). Nevertheless, the total time complexity of calculating the *p*-value is definitely dominated by the compexity of applying DP algorithm to each chromosome to compute all *N*_*q*_(*k*_*q*_). As DP algorithm on each chromosome is done independently, we can take advantage of parallel computing and the total running time would be 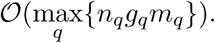

---

**Figure S1:**
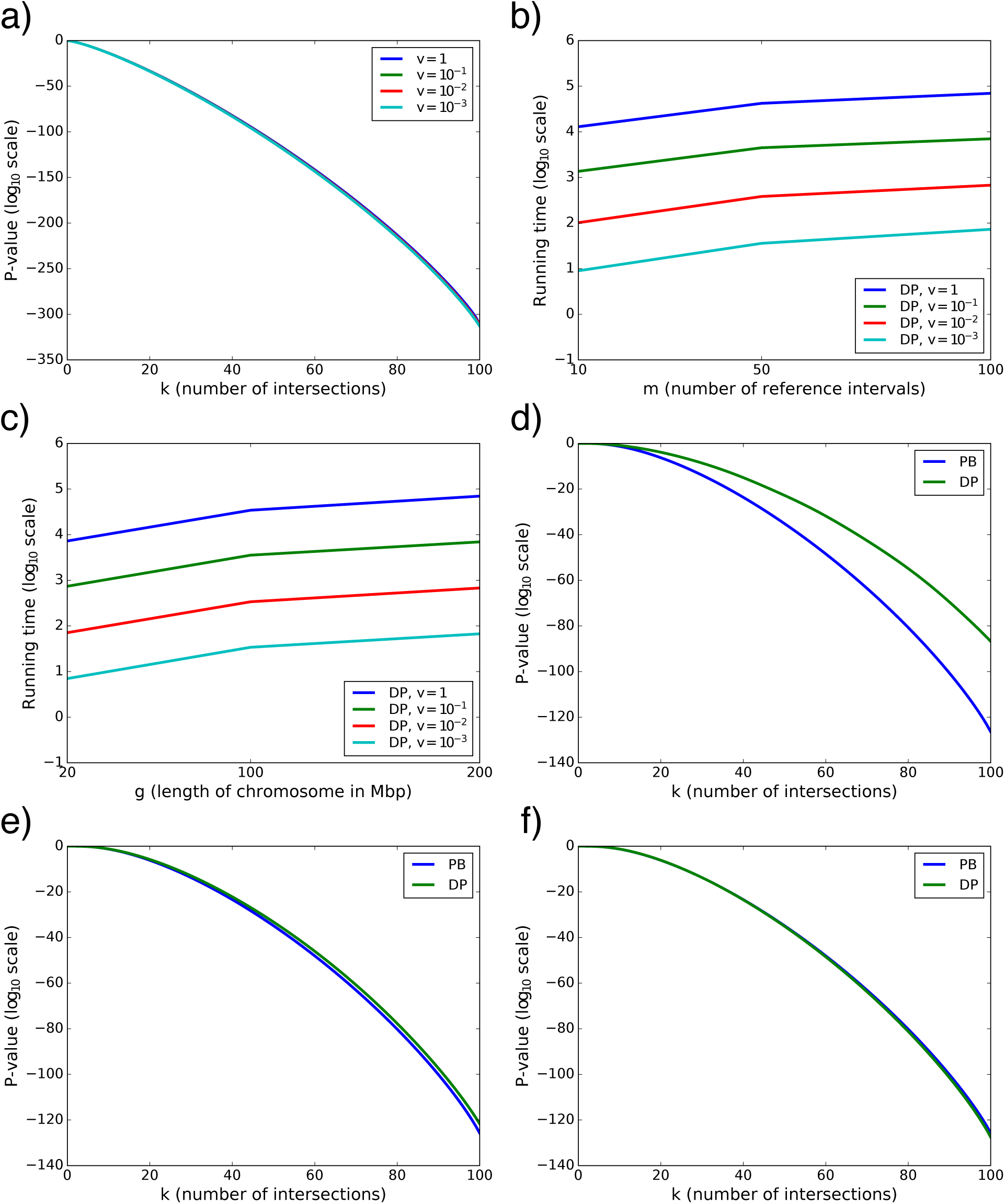
Further evaluation of methods using simulated datasets. (a) Impact of scaling parameter *ν* on DP *p*-value when *l*_*i*_, *x*_*j*_ ~ 𝒖[100bp, 4Kbp]. (b) Running time (secs.) of DP algorithm as a function of *m*, with *n* = 100, *g* = 200Mbp. (c) Running time (secs.) of DP algorithm as a function of *g*, with *n* = *m* = 100. (d– f) Impact of approximation on *p*-value computation. Simulations are run with *g* = 200Mbp, *m* = 100, *n* = 1000, *l*_*i*_, *x*_*j*_ ~ 𝒖[1Kbp, 2Kbp]; (d) *η* = 0.0079. (e) *η* = 0.062. (f) *η* = 0.68.

